# Building a better biofilm - formation of *in vivo*-like biofilm structures by *Pseudomonas aeruginosa* in a porcine model of cystic fibrosis lung infection

**DOI:** 10.1101/858597

**Authors:** Niamh E. Harrington, Esther Sweeney, Freya Harrison

## Abstract

*Pseudomonas aeruginosa* biofilm infections in the cystic fibrosis (CF) lung are highly resistant to current antimicrobial treatments and are associated with increased mortality rates. The existing models for such infections are not able to reliably mimic the clinical biofilms observed. We aimed to further optimise an *ex vivo* pig lung (EVPL) model for *P. aeruginosa* CF lung infection that can be used to increase understanding of chronic CF biofilm infection. The EVPL model will facilitate discovery of novel infection prevention methods and treatments, and enhanced exploration of biofilm architecture. We investigated purine metabolism and biofilm formation in the model using transposon insertion mutants in *P. aeruginosa* PA14 for key genes: *purD*, *gacA* and *pelA*. Our results demonstrate that EVPL recapitulates a key aspect of *in vivo P. aeruginosa* infection metabolism, and that the pathogen forms a biofilm with a clinically realistic structure not seen in other *in vitro* studies. Two pathways known to be required for *in vivo* biofilm infection - the Gac regulatory pathway and production of the Pel exopolysaccharide - are essential to the formation of this mature, structured biofilm on EVPL tissue. We propose the high-throughput EVPL model as a validated biofilm platform to bridge the gap between *in vitro* and CF lung infection.

## Introduction

Biofilms are microbial aggregates that consist of bacterial cells and a self-produced matrix primarily composed of proteins, lipids, polysaccharides and nucleic acids [1]. The biofilm matrix provides the microbial population with increased antibiotic tolerance and protection against the host immune response [2]. The persistent, biofilm-associated lung infections that affect people with the genetic condition Cystic Fibrosis (CF) are highly resistant to antibiotic treatment and, despite being well studied, continue to have a large impact on health and life expectancy. More than 70,000 people worldwide are believed to have CF, though it is thought to be widely underdiagnosed across Asia as epidemiological reports are largely derived from data collected in western countries [3–5]. Despite the rising average life expectancy of CF patients [6], it still has one of the highest mortality rates among human genetic conditions [5].

CF lung infections are polymicrobial [7], however *Pseudomonas aeruginosa* is the most abundant CF pathogen in the UK. It is estimated that 75% of adults with CF are either chronically or transiently infected with *P. aeruginosa* during their lifetime [8, 9]. By 5 years old approximately 53% of children with CF are infected with *P. aeruginosa* [10, 11]. Detection of *P. aeruginosa* infection is associated with worsened disease outcomes including rapid lung function decline and higher mortality rates [12–14].

*P. aeruginosa* is able to switch from the initial acute stage of infection, where virulence is high, infection is transient and biofilms are not formed [15], to chronic biofilm infection. This chronic infection is characterised by embedding of *P. aeruginosa* biofilm aggregates in the mucus plugs typically found in CF individuals’ airways [16]. Chronic infection is traditionally difficult to replicate *in vitro* and so the ability to study *P. aeruginosa* virulence, persistence and treatment response at this stage of infection is limited. Once established, chronic *P. aeruginosa* infection is almost impossible to eradicate, and there is a poor correlation between antibiotic susceptibility testing and the clinical outcomes of antibiotic treatment [17, 18]. This is largely a result of altered resistance phenotypes in *in vivo* biofilms compared to *in vitro* [19]. Without the development of novel models to study clinically realistic, CF-associated chronic biofilm infections in the laboratory, development of more effective treatments to prevent or treat these infections will continue to be hindered.

There are a number of models currently used to study *P. aeruginosa* in mono- and polymicrobial CF biofilm infections. These include, but are not limited to, artificial sputum media (ASM) [20–23], *in vitro* surface-attached biofilm models [24] and murine models with cystic fibrosis transmembrane conductance regulator mutations that mimic the mutations causative of CF in humans [25]. Limitations with such model systems have been identified. These include the distinct difference in *P. aeruginosa* transcriptomic profiles between isolates grown in ASM and direct transcriptomic analysis of CF sputum samples [26]. The structure of biofilms grown in *in vitro* systems also differ from the biofilms observed *in vivo* [27]. In addition, mouse models are unable to reliably mimic the *P. aeruginosa* infections that occur in people with CF [28]. The airway secretions from CF mice have been shown to be very different from human airway secretions [29], which could provide explanation for the significant differences in *P. aeruginosa* transcriptome found between mouse and human CF infection [26].

We previously published brief details of a novel *ex vivo* pig lung (EVPL) model for *P. aeruginosa* biofilm infection of CF bronchioles. The model combines one formulation of ASM with sections of porcine bronchiole [30]. The EVPL bronchiolar tissue demonstrated potential in recapitulating clinically relevant chronic infection [30, 31]. In comparison to more commonly used model organisms, including rats and mice, pig lungs demonstrate closer similarity to human lungs. The two are similar in their metabolic composition and overall physiology, anatomy and immunology [29, 32]. The lungs used for the model are obtained from pigs slaughtered for meat, thus the EVPL model offers a replacement for live animal (rodent) infection models, which is clinically realistic, economical and ethical.

Here, we provide detailed validation of EVPL as a model for chronic *P. aeruginosa* biofilm infection in CF. We have built on our previous work to demonstrate that the model is able to successfully mimic clinically relevant aspects of *in vivo* CF *P. aeruginosa* metabolism and biofilm formation. We show key pathways required for CF *P. aeruginosa* biofilm formation and maturation *in vivo* are similarly essential in EVPL. Our findings also highlight the importance of visualising biofilms in order to understand chronic infection in a way that *in vitro* models of bacterial growth cannot achieve. EVPL has the potential to provide a platform to aid in the development of novel drugs and treatment regimens, which can be used to target the growing incidence of highly antibiotic-tolerant biofilms in the unique environment of CF lungs.

## Materials and Methods

### Bacterial strains

Two isolates of *P. aeruginosa* (SED42, SED43), from a chronically-colonised person with CF, were used as example clinical isolates. These strains have previously been phenotypically characterised [33]. *P. aeruginosa* PA14 transposon insertion mutant strains (Table 1) and the isogenic wild type (WT) used in this work were obtained from the PA14 Non-Redundant Transposon Insertion Mutant Set [34], [35]. The transposon’s presence in the correct locus of the putative PA14 mutants for *leuB*, *purD*, *gacA* and *pelA* was confirmed by arbitrary PCR and Sanger sequencing [34], [35]. No sequence was obtained from three repeated PCRs for the *metX* mutant hence locus-specific primers for *metX* were designed (5’-ATCGGGTCCTCACACGCTG-3’ and 5’-TCCGACCCTGAGTTCCTCG-3’). PCR and gel electrophoresis confirmed amplification of a fragment in the putative *metX* mutant of ∼1.5 kb. This is consistent with the presence of the transposon (994 bp); the expected fragment in the absence of a transposon is ∼500 bp.

**Table 1.** Description of the *Pseudomonas aeruginosa* PA14 transposon insertion mutants used for our work [34, 35]. Information for each gene is provided and the associated CF infection phenotype following loss of gene function [23,36,37].

| Mutant ID | Active Gene ID | Active Gene Name | Active Gene Description | Predicted Infection Phenotype in EVPL |
| --- | --- | --- | --- | --- |
| 31800 | GID1896 | *leuB* | 3-isopropylmalate dehydrogenase | No effect |
| 23405 | GID1574 | *metX* | Homoserine O-acetyltransferase | No effect |
| 29794 | GID1301 | *purD* | Phosphoribosylamine-glycine ligase | No effect |
| 54630 | GID3719 | *gacA* | Response regulator GacA | Biofilm maturation defects |
| 26187 | GID86 | *pelA* | Conserved hypothetical protein | Unable to form biofilm |

### Artificial sputum media

The first artificial sputum medium (ASM) version, ASM1, was prepared following a published recipe, also referred to as SCFM [22]. The second ASM version: ASM2, is a slightly modified version of ASM1 and was similarly prepared following a published method, also referred to as SCFM2 [23]. For instances where glucose was removed from these recipes, it was substituted with an equivalent volume of deionised water.

### *In vitro P. aeruginosa* growth and biofilm formation in ASM

Colonies of PA14, SED42 and SED43 were taken from Luria-Bertani (LB) agar plates (Melford) and grown overnight at 37 °C. Colonies were used to inoculate replicate 100 µl aliquots of ASM1 or ASM2, ± 3 µM glucose, in a 96-well microplate with a peg lid (Innovotech), n = 4 per growth condition. Cultures were incubated at 37 °C for 24 h. Growth of planktonic fractions of the populations was assessed by briefly shaking the well plate then reading absorbance at 600 nm using a Tecan Spark 10M plate reader. Biofilm formation on pegs was quantified using a crystal violet assay [38]. Briefly, pegs were rinsed with 150 µl phosphate-buffered saline (PBS) in a fresh 96-well plate to remove loosely-adhering cells. The peg lid was then transferred to a further 96-well plate with 150 µl 0.1% (w/v) crystal violet (Vickers Laboratories) per well. It was incubated at room temperature for 15 min and the pegs rinsed twice in PBS as above, before drying in a laminar flow hood for 30 min. The pegs were transferred to another 96-well plate containing 150 µl 95% (v/v) ethanol per well to solubilise the crystal violet. Absorbance of the solubilised crystal violet was read at 590 nm in a Tecan Spark 10M. The relative allocation of bacterial cells to the biofilm per well was calculated by dividing A590nm of crystal violet by A600nm of corresponding planktonic subpopulation.

### Microscopy of *P. aeruginosa* grown *in vitro* in ASM

Three replicate 100 µl aliquots of ASM1 or ASM2, ± 3 µM glucose, per growth condition were inoculated with colonies of PA14, SED42 or SED43 taken from LB agar plates grown overnight at 37 °C. Cultures were incubated at 37 °C for 36 h. Aliquots were taken of each culture, diluted 1:100 in PBS and 10 µl heat fixed on a glass slide. Each fixed, diluted culture was then Gram stained. The stained slides were imaged using a Zeiss Axio Scope.A1 light microscope with the Zeiss Axiocam Erc 5s and Zeiss Zen 2.3 pro software.

### *In vitro* growth of *P. aeruginosa* metabolic mutants

Cultures were grown for 6 h in LB at 37 °C. Bacterial cells were washed and starved in PBS for 2 h, then resuspended in ASM1 or ASM2. One set of ASM cultures were left as per the original method, and the other supplemented with either 20 µg ml^-1^ leucine, 20 µg ml^-1^ methionine or 30 µg ml^-1^ each of guanine, adenine, xanthine and hypoxanthine as relevant for each mutant. Purines had to be dissolved in NaOH, so the comparative original ASM was supplemented with an equivalent amount of NaOH. Subsequently, 150 µl of each resuspended culture was transferred to a 96-well plate and grown for a further 24 h at 37 °C in a Tecan Spark 10M. Every 20 min, the plate was shaken for 5 s and the absorbance read at 600 nm.

### EVPL dissection and infection

EVPL was prepared as previously described [30]. Briefly, porcine lungs were obtained from a local butcher (Quigley and Sons, Cubbington) and dissected on the day of delivery under sterile conditions. The pleura of the ventral surface was heat sterilised using a hot pallet knife. A sterile razor blade was then used to make an incision in the lung, exposing the bronchiole. A section of the bronchiole was extracted and the exterior alveolar tissue removed using dissection scissors. Bronchiolar sections were washed once in a 1:1 mix of Dulbecco’s modified Eagle medium (DMEM) and RPMI 1640 supplemented with 50 µg ml^-1^ ampicillin (Sigma-Aldrich) then cut into ∼5 mm wide longitudinal strips. The bronchiolar strips were placed in a second 1:1 DMEM, RPMI 1640 + 50 µg ml^-1^ ampicillin wash and cut into squares (∼5 x 5 mm). The tissue squares were washed for a third time in 1:1 DMEM, RPMI 1640 + 50 µg ml^-1^ ampicillin. Bronchiolar pieces were then further washed in ASM, UV sterilised for 5 min and transferred to individual wells of a 24-well plate containing 400 µl ASM (+ 20 µg ml^-1^ ampicillin) solidified with 0.8% (w/v) agarose per well.

A sterile 29G hypodermic needle (Becton Dickinson Medical) was touched to the surface of a *P. aeruginosa* colony grown on LB agar overnight at 37 °C and used to pierce an individual piece of bronchiolar tissue. Uninfected control tissue sections were mock inoculated with a fresh, sterile needle. Following infection of all tissue pieces, 500 µl of ASM + 20 µg ml^-1^ ampicillin was added to each well. Tissue was incubated at 37 °C with a Breathe-Easier^®^ membrane (Diversified Biotech) for the desired length of time.

### EVPL biofilm recovery and assessment of bacterial load

Bronchiolar tissue pieces were removed from the 24-well plate following incubation, and each briefly washed in 500 µl PBS in a fresh 24-well plate. Tissue pieces were then transferred into sterile homogenisation tubes (Fisherbrand) containing eighteen 2.38 mm metal beads (Fisherbrand) and 1 ml PBS. Tissue was bead beaten in a FastPrep-24 5G (MP Biomedicals) for 40 s at 4 m s^-1^ to recover the bacteria and virulence factors from the tissue-associated biofilm. To determine the bacterial load, the homogenate was serially diluted in PBS and plated on LB agar. Plates were incubated overnight at 37 °C and colony counts used to calculate colony forming units (CFU) per tissue piece.

### Determination of *P. aeruginosa* virulence factor production in EVPL biofilms

To quantify virulence factors produced by the *P. aeruginosa* biofilm population, the homogenate was diluted 1:4 in PBS to obtain a sufficient volume for further experiments. The diluted homogenate was passed through a 0.2 µm filter to remove bacterial cells and tissue debris.

To assay the amount of the siderophores pyoverdine and pyochelin present in homogenates, 100 µl aliquots of diluted sterile homogenate were transferred to a black 96-well plate. Fluorescence was measured with excitation/emission of 400 ± 20/460 ± 20 nm for pyoverdine [39] and 360 ± 35/460 ± 20 nm for pyochelin [40] in a Tecan Spark 10M.

To measure total protease, 100 µl of diluted sterile homogenate was mixed with 5 mg azocasein (Sigma-Aldrich) dissolved in 900 µl 100 mM Tris-HCl + 1 mM CaCl2. The reaction mixture was incubated with 170 rpm shaking for 2 h at 37 °C, then stopped by the addition of 100 µl 120 mM EDTA. The mixture was centrifuged at 13,000 rpm for 1 min and the absorbance of the supernatant read at 400 nm.

The amount of two quorum sensing (QS) signals in the diluted sterile homogenate was assayed using *Escherichia coli* biosensors [41]. Expression of a *luxCDABE* reporter in response to either 3-oxo-C12-HSL or C4-HSL was measured as relative light units (RLU) for *E. coli* strains carrying the reporter plasmids pSB1075 or pSB406 respectively.

### Extracellular DNA detection in EVPL biofilms

Following 2 d and 7 d post-infection (PI), homogenate from EVPL cultures of PA14 and SED43 and uninfected control tissue were centrifuged at 13,000 rpm for 1 min (n = 3). The pellet was run through a Promega Wizard® Genomic DNA extraction kit. Recovered DNA was eluted in 50 µl sterile water and 10 µl of the eluate was loaded onto a 0.7% (w/v) agarose + SYBR™ Safe stain (Invitrogen) gel. Following electrophoresis, the resulting gel was visualised under blue light.

### Crystal violet assay on EVPL biofilms

Infected and uninfected control bronchiolar tissue pieces were placed in a 24-well plate with 500 µl PBS per well. The plate was briefly shaken to remove any loosely-adhering *P. aeruginosa* cells from the tissue. Tissue pieces were then moved to another 24-well plate with 500 µl 0.1% (v/v) crystal violet per well, and incubated at room temperature for 15 min with 170 rpm shaking. Tissue sections were transferred into a fresh 24-well plate and the crystal violet treatment and incubation was repeated. The tissue was then transferred to a further 24-well plate and left to dry in a laminar flow hood for 30 min. The tissue and biofilm bound crystal violet was solubilised in 500 µl 95% (v/v) ethanol and incubated at room temperature for 15 min with 170 rpm shaking. The tissue was removed from the wells and absorbance read at 590 nm in a Tecan Spark 10M.

### Hematoxylin & eosin staining

*P. aeruginosa* infected EVPL tissue pieces and uninfected control tissue were fixed in 10% (v/v) neutral buffered formalin (VWR Chemicals). The tissue samples were sent to the University of Manchester’s Histology Core Facility for paraffin wax embedding, sectioning and mounting. Mounted tissue sections were de-paraffinized in xylene for 20 min. To re-hydrate the tissue, slides were transferred to 95% (v/v) ethanol followed by 70% (v/v) ethanol. Any residual ethanol was removed by washing slides in distilled water. Samples were stained in Mayer’s hemalum solution (Merck Millipore) then washed in running ‘tap water’ (1 L tap water with 2 tsp sodium carbonate). Samples were counterstained in eosin Y solution (Merck Millipore) for then dehydrated by dipping the slides in 95% (v/v) ethanol. Samples were transferred to fresh 95% (v/v) ethanol then placed in 100% (v/v). The samples were then placed in xylene. The samples were mounted using DPX mounting fluid and images taken using a Zeiss Axio Scope.A1 light microscope with the Zeiss Axiocam Erc 5s and Zeiss Zen 2.3 pro software.

### Gram staining

*P. aeruginosa* infected EVPL tissue pieces, plus uninfected controls, were fixed and paraffin-embedded as described above for H & E staining. The tissue sections were de-paraffinized in xylene. Samples were re-hydrated in 100% (v/v) ethanol for then moved to fresh 100% (v/v) ethanol. Slides were placed in 95% (v/v) isopropanol then 70% (v/v) isopropanol. Residual alcohol was removed by washing slides in distilled water. Crystal violet (Pro-Lab Diagnostics) was applied to the samples and rinsed with water, then iodine (Pro-Lab Diagnostics) was applied. The iodine was then rinsed from the samples using tap water and acetone added for decolourisation. Tap water was used to rinse off the acetone and 1% (v/v) neutral red (Pro-Lab Diagnostics) applied to counterstain. The slides were again rinsed with tap water and blotted dry with filter paper. The samples were dehydrated in 100% (v/v) isopropanol then placed in two fresh changes of 100% (v/v) xylene. Samples were mounted using DPX mounting fluid and images taken using a Zeiss Axio Scope.A1 light microscope with the Zeiss Axiocam Erc 5s and Zeiss Zen 2.3 pro software.

### Statistical analysis

Data were analysed using either R v3.5.1 or RStudio (v1.2.1335 on Mac OS Mojave v10.14.6) [42], with *P* < 0.05 considered significant. The *car* package [43] was used to conduct ANOVAs with type II SS when missing values caused non-orthogonality (i.e. analyses of C4-HSL and total protease). The *multcomp* package [44] was used for post-hoc comparisons. The *FactoMineR* package [45] was used for principal components analysis. All raw data and R code used for analyses, including data transformations used to meet model assumptions, are provided in the Data Supplement.

## Results

### Simple and complex formulations of artificial CF sputum lead to comparable *P. aeruginosa* growth and virulence in EVPL

Various formulations of artificial CF sputum media have been published [20–23]. Here we focus on two formulations, which we term ASM1 and ASM2. ASM1 is the original formula we chose for use with the EVPL model [30,31,46,47] and ASM2 is an updated version that has since been published by the same laboratory [23]. ASM1 contains concentrations of free amino acids, cations, anions and lactate that are representative of the average concentrations found in a selection of sputum samples from people with CF. ASM1 was initially chosen for use with EVPL as it was shown to cue comparable carbon-usage pathways and expression of quorum sensing signals by *P. aeruginosa* PA14 to growth in medium made from lyophilised patient sputum [22]. ASM2 adds bovine maxillary mucin, membrane lipid 1,2-Dioleoyl-sn-glycero-3-phosphocholine (DOPC), N-acetyl glucosamine (NAG) and free DNA. These additions were made to better represent the presence of host mucins, bacterial membrane and wall components, and the extracellular DNA (eDNA) found in the biofilm plugs formed in the bronchioles of people with CF. ASM2 has been shown to result in a comparable essential genome for *P. aeruginosa* PA14 to lyophilised patient sputum media *in vitro* [23]. We aimed to determine whether the EVPL model required the more complex ASM2, or whether EVPL + ASM1 would provide a sufficient platform for laboratory growth of *P. aeruginosa* with more clinically realistic characteristics.

In our previous work with EVPL, we removed glucose from the original ASM1 formulation as it appeared to enhance the growth of endogenous bacteria from the porcine lungs [31]. We have tested the effects of glucose removal from both versions of ASM on *P. aeruginosa* growth and biofilm formation *in vitro* (Figure S1). Our results show small and inconsistent effects of ASM version and glucose removal on PA14 and two clinical CF isolates. The limited effect of glucose removal is consistent with the expectation that *P. aeruginosa* growth in the CF lung environment mostly depends on using amino acids and short-chain fatty acids as carbon sources [22,23,48,49]. Hence ASM without glucose is the optimal approach for use with EVPL, so glucose was omitted from ASM for all further work.

When grown for 36 h *in vitro*, *P. aeruginosa* appeared to form biofilm-like aggregates in ASM2 that were not observed in ASM1, where bacterial cells appeared stressed (Figure S2). However our previous work showed morphologically ‘normal’ *P. aeruginosa* growing as extensive biofilms on EVPL + ASM1 4 d PI [30]. This suggests the presence of EVPL overcomes these differences in biofilm formation; therefore, we investigated the effect of ASM version on growth and virulence in EVPL. *P. aeruginosa* PA14 and clinical isolates SED42 and SED43 were used to infect EVPL with either ASM1 or ASM2, and biofilm was recovered 2 d post-infection (PI). The biofilm load did not differ between EVPL + ASM1 and EVPL + ASM2 for PA14 or the exemplar clinical isolates (Figure 1A). To confirm that ASM version did not impact on other aspects of *P. aeruginosa* biology, we measured the production of a selection of exoproducts associated with virulence: the quorum sensing signals C4-HSL and 3-oxo-C12 HSL, total protease and the siderophores pyoverdine and pyochelin (Figure 1B). ANOVAs confirmed that there was not a significant effect of ASM version on any of these variables, as either a main effect or interaction with strain (Table 2). Principal component analysis did not reveal any clear clustering of phenotypes by ASM version (Figure S3). We also found that EVPL + ASM1 was able to maintain extended *P. aeruginosa* growth over 14 d (Figure S4), with detectable amounts of eDNA present in the biofilms formed 7 d PI (Figure S5). These results indicate the addition of EVPL removes the requirement of further additions made to ASM1 in the form of ASM2, and enhances the suitability of the simpler artificial media to mimic CF infection.

**Figure 1.**
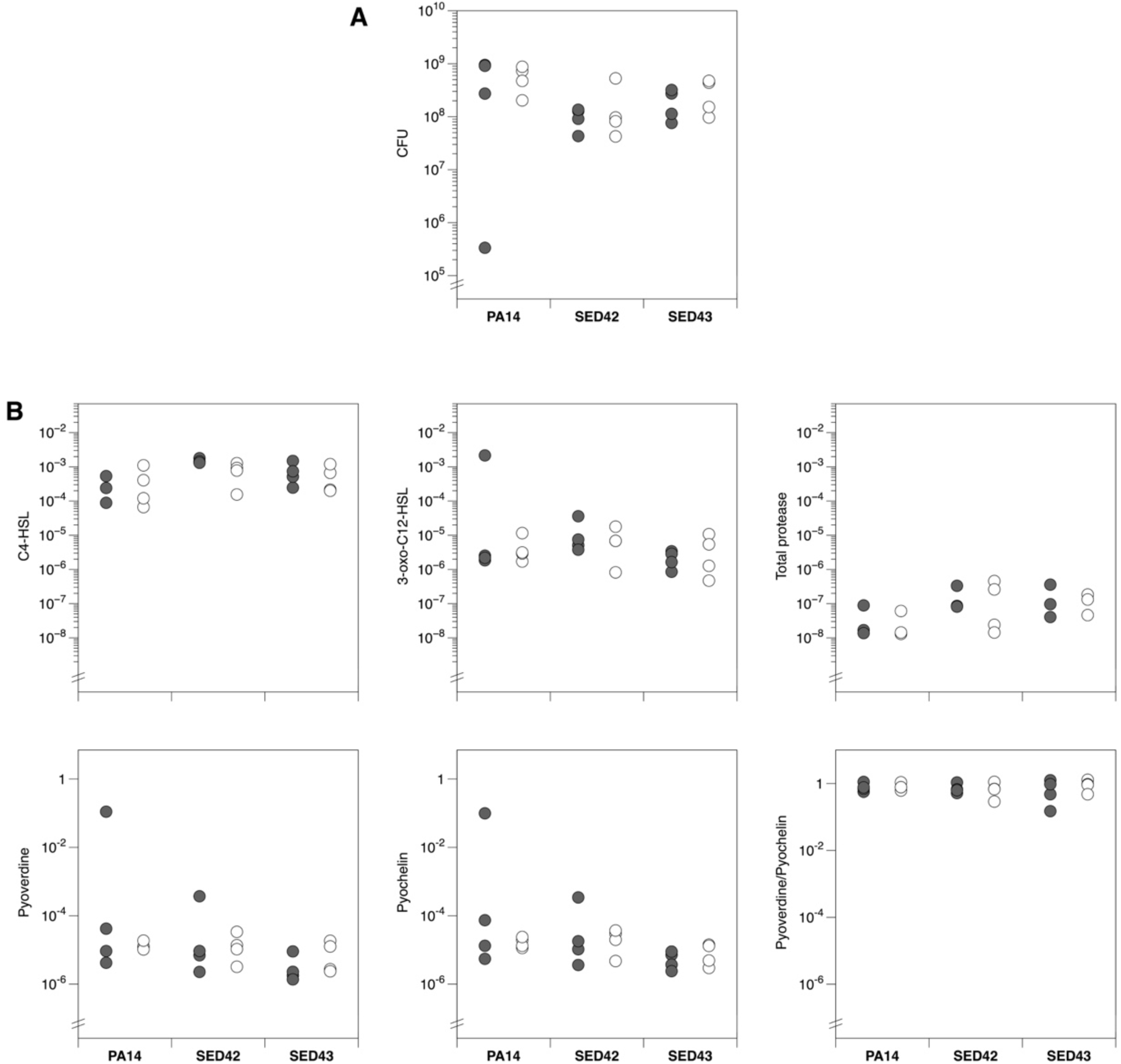
(A) Growth of the laboratory isolate *Pseudomonas aeruginosa* PA14 and two exemplar *P. aeruginosa* Cystic Fibrosis (CF) isolates: SED42 and SED43, on four pieces of *ex vivo* pig lung (EVPL) bronchiole plus either artificial sputum media 1 (ASM1) (closed circles) or ASM2 (open circles). The colony forming units (CFU) were retrieved from biofilm after 2 d growth. Strains varied in their biofilm load (ANOVA: F_1,18_ = 3.64, *p* = 0.047), but it was not affected by ASM version (main effect F_1,18_ = 0.84, *p* = 0.37; interaction with strain F_2,18_ = 0.01, *p* = 0.99). Results were unaffected if the outlier in PA14 + ASM1 was excluded (*p*-values of 0.002, 0.70 and 0.64, respectively). (B) Production of selected virulence factors by *P. aeruginosa* isolates in EVPL + ASM1 (closed circles) or EVPL + ASM2 (open circles). Units are as follows: for C4-HSL and 3-oxo-C12-HSL, Relative Light Units (RLU) of *Escherichia coli* biosensor per CFU of *P. aeruginosa*; for total protease, A_400_ following azocasein assay per CFU of *P. aeruginosa*; for pyoverdine and pyochelin, fluorescence per CFU of *P. aeruginosa* individually and as a ratio.

**Table 2.** Results of ANOVAs for the data shown in Figure 1B. None of the analyses show a significant effect of ASM version, either as a main effect or as an interaction with strain. Raw data and R code are provided in the Data Supplement.

|  | Strain |  | ASM |  | Strain*ASM |  |
| --- | --- | --- | --- | --- | --- | --- |
|  | F<sub>2,18</sub> | *p* | F<sub>1,18</sub> | *p* | F<sub>2,18</sub> | *p* |
| **C4-HSL** | 6.00 | 0.011* | 2.27 | 0.150 | 1.86 | 0.185 |
| **3-oxo-C12-HSL** | 1.41 | 0.342 | 0.49 | 0.493 | 0.35 | 0.709 |
| **Total protease** | 2.07 | 0.155 | 0.37 | 0.553 | 0.95 | 0.406 |
| **Pyoverdine (PVD)** | 2.07 | 0.155 | 0.37 | 0.553 | 0.95 | 0.406 |
| **Pyochelin (PCH)** | 2.14 | 0.147 | 0.65 | 0.430 | 0.89 | 0.429 |
| **PVD/PCH** | 0.25 | 0.781 | 0.26 | 0.619 | 0.28 | 0.758 |

### EVPL removes the essentiality of purine biosynthesis for *P. aeruginosa* growth in both simple and complex formulations of ASM

To further confirm the optimal ASM version for use in all subsequent work, and the effect on *P. aeruginosa* of adding EVPL tissue to ASM, we compared one key aspect of primary metabolism in EVPL + ASM1 and EVPL + ASM2. Purine biosynthesis was selected as it was previously found to be essential in ASM2 *in vitro* but not in sputum taken from people with CF [23]. This lack of consistency with true infection highlights a specific area where the composition of ASM2 appears sub-optimal as a model for CF.

To confirm this previous finding, we performed *in vitro* growth curves in ASM1 and ASM2 of the laboratory strain wild type (WT) *P. aeruginosa* PA14 and a PA14 transposon insertion mutant for the purine auxotroph gene *purD*. We also measured growth of two isogenic mutants auxotrophic for amino acids abundant in ASM: *leuB* and *metX*, as positive controls. We found that in both versions of ASM purine supplementation is required for *purD* mutant growth *in vitro* (Figure 2A).

**Figure 2.**
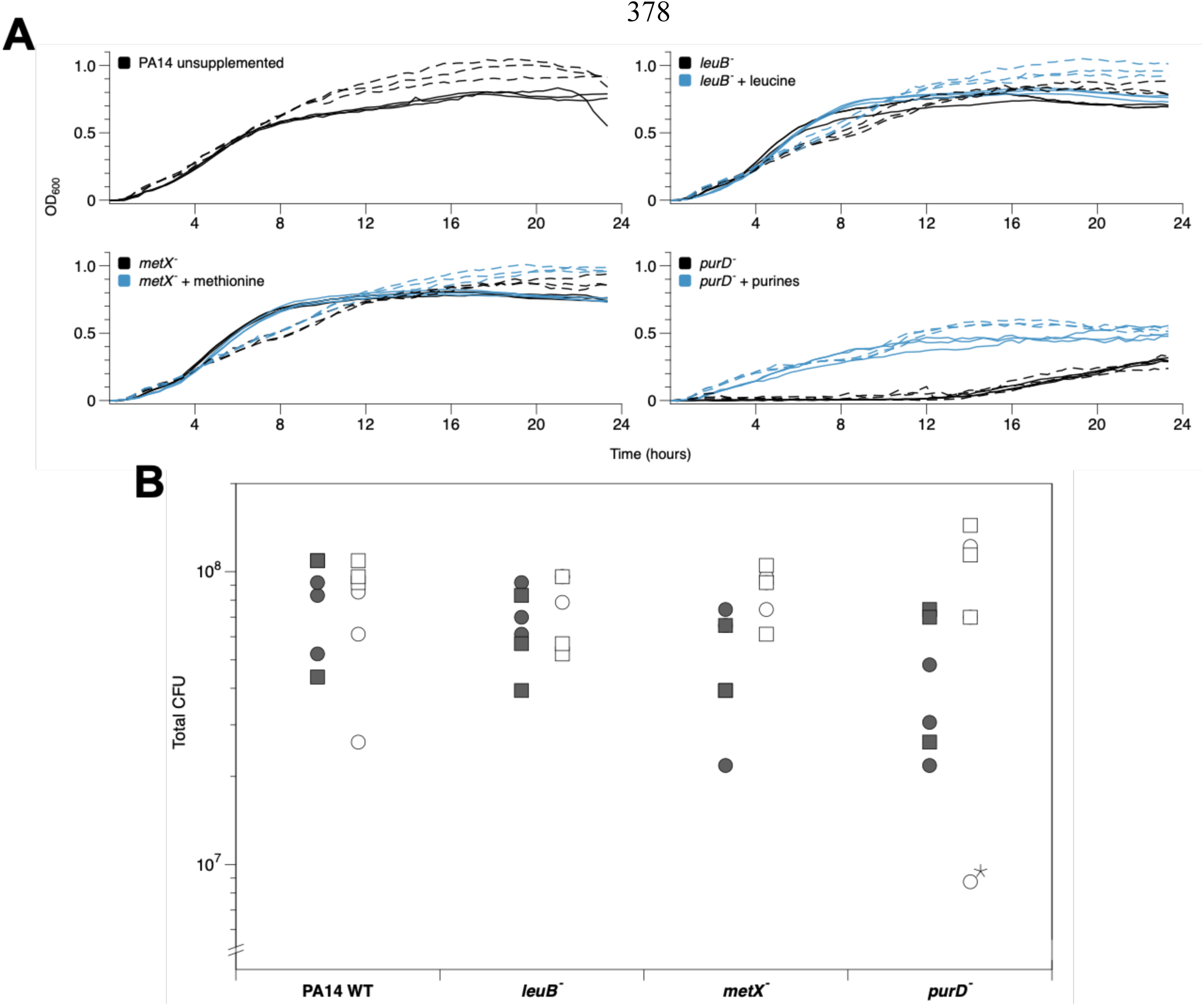
(A) Growth of *Pseudomonas aeruginosa* PA14 and transposon insertion metabolic mutants *in vitro* in artificial sputum media 1 (ASM1) (solid) and ASM2 (dashed). Black lines are ASM alone, blue lines show ASM cultures supplemented with 20 µg ml^-1^ leucine, 20 µg ml^-1^ methionine or 30 µg ml^-1^ each of purines (guanine, adenine, xanthine and hypoxanthine) as appropriate. All treatments were replicated in triplicate and lines show individual replica cultures. (B) Growth of *P. aeruginosa* PA14 and transposon insertion metabolic mutants on three pieces of *ex vivo* pig lung (EVPL) bronchiole from each of two independent lungs (circles vs. squares) plus ASM1 (closed symbols) or ASM2 (open symbols). The graph shows colony-forming units (CFU) retrieved from individual EVPL biofilms after 48 h growth. An ANOVA was run to test for effects of lung identity, genotype, version of ASM used and the interaction between genotype and version of ASM used, on biofilm load. One clear outlier (*purD^-^*, ASM2, denoted with a star) was identified and excluded from analysis of this data. There was no main effect of lung identity or genotype on biofilm load (ANOVA: F_3,38_ = 0.56, p = 0.448 and F_1,38_ = 0.46, p = 0.712). Biofilm load was affected by the version of ASM used (main effect F_1,38_ = 12.1, p = 0.001; interaction with genotype F_3,38_ = 3.50, p = 0.024). A Tukey test showed statistical interaction was driven by the *purD* mutant (p < 0.01). Planned contrasts by conducting t-tests where t = (mutant CFU – WT CFU in the same medium)/SE_diff_, where SE_diff_ is the standard error of the difference, calculated using the residual mean square from the ANOVA, showed no significant difference in CFU between individual mutants and WT in the same version of ASM (2-tailed p ≥ 0.126).

To identify whether the presence of EVPL could compensate for this shortcoming we infected two independent lungs with the four *P. aeruginosa* PA14 strains used *in vitro* (WT; *purD, leuB* and *metX* mutants). Bacterial growth was recovered at 2 d PI to determine CFU per lung (Figure 2B). This was repeated using either ASM1 or ASM2 with the EVPL. Figure 2B highlights the consistency in CFU per lung of the WT between EVPL + ASM1 and EVPL + ASM2. Statistical analysis found no significant difference in CFU between any individual mutant and the WT in the same version of ASM (Figure 2B). Crucially, the *purD* mutant demonstrated no reduction in bacterial load compared with the WT in the corresponding ASM version. It can be concluded that addition of EVPL removes the essentiality of purine biosynthesis in ASM1, making it consistent with clinical infection. It is also worth noting that the slight growth defect of the WT observed in ASM1 *in vitro* compared to ASM2 (Figure 2A) is not observable when EVPL is added (Figure 2B).

In conclusion, there does not appear to be any significant differences in *P. aeruginosa* growth, virulence or purine metabolism in EVPL + ASM1 and EVPL + ASM2. We therefore used EVPL + ASM1 to explore the architecture of *P. aeruginosa* biofilms formed.

### *P. aeruginosa* biofilm formation on EVPL with ASM recapitulates the morphology of *in vivo* CF biofilms, and is dependent on the Gac regulatory system and Pel polysaccharide

Biofilm formation is arguably the defining phenotype of chronic *P. aeruginosa* lung infection in individuals with CF. Following identification of optimal EVPL model conditions, we explored the structure and mechanisms of biofilm formation by *P. aeruginosa* PA14 on EVPL + ASM1. We compared the biofilm formed by WT PA14 with those formed by transposon insertion mutants for the genes *gacA* and *pelA*. Both *gacA* and *pelA* are important in biofilm infection by *P. aeruginosa* PA14; either by direct regulation (*gacA*) or production of a structural matrix polysaccharide (*pelA*). GacA is a response regulator from the global activator of antibiotic and cyanide synthesis (Gac) two-component regulatory system. The Gac system is well-studied and known to be largely responsible for the switch from acute to chronic infection; in particular it regulates biofilm maturation [50, 51]. PelA is a periplasmic protein required for biosynthesis of the biofilm matrix exopolysaccharide Pel. It is one of three exopolysaccharides associated with *P. aeruginosa* biofilm formation: Pel, Psl and alginate. In PA14, alginate is non-essential for biofilm formation and PA14 is Psl deficient. Hence, biofilm formation in this strain requires the Pel exopolysaccharide [52].

To measure and visualise the biofilm formed by *P. aeruginosa* PA14 WT and the *gacA* and *pelA* mutants, strains were inoculated individually into replica lung pieces from three independent lungs. Uninfected tissue was used as the negative control. We recovered the biofilm at 2 d and 7 d PI to determine CFU per EVPL section across an extended time period (Figure 3A). ANOVAs identified a significant difference in CFU per EVPL tissue section between both the *gacA* and *pelA* mutants and the WT at 2 d PI (Figure 3A). However, there was only a significant difference in CFU per EVPL section between the *gacA* mutant and WT 7 d PI (Figure 3A). The interaction term between replicate lungs and infection strain was found to be significant, reflecting the natural variation between lungs. This is not dissimilar to the patient-to-patient variation expected clinically.

**Figure 3.**
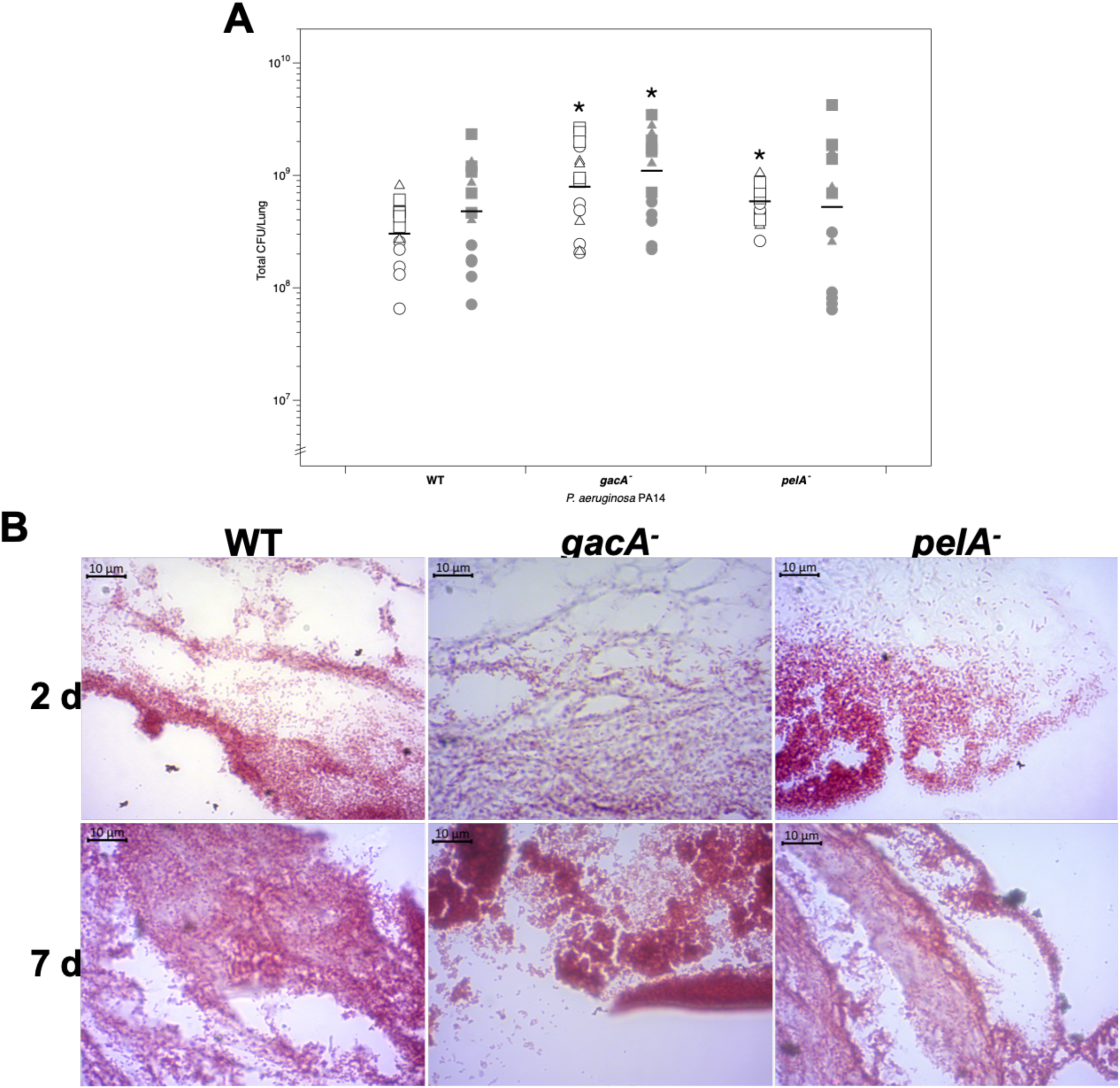
(A) Colony forming units (CFU) per *ex vivo* pig lung (EVPL) tissue section infected with *Pseudomonas aeruginosa* PA14 wild type (WT) and transposon mutants at 2 d (open symbols) and 7 d (closed symbols) post-infection (PI). Five replicate pieces of tissue were infected per strain from three independent lungs. Bars denote means, and asterisks denote a significant difference from the WT under that condition (post-hoc Dunnett’s test after ANOVA). ANOVA found a significant difference between strains, lung and their interaction at 2 d PI (strain F_4,60_ = 10.27, P < 0.01; lung F_2,60_ = 26.05, P < 0.01; interaction F_8,60_ = 1.99, P = 0.06). Post-hoc analysis showed that there was significantly higher CFU recovered of the *gacA* mutant compared to the WT (P < 0.01), and higher CFU of *pelA* mutant than the WT (P = 0.01). At 7 d PI a significant difference in bacterial recovery was found between strains and between lungs, as well as a significant interaction between strain and lung (ANOVA: strain F_4,60_ = 5.40, P < 0.01; lung F_2,60_ = 78.98, P < 0.01; interaction F_8,60_ = 2.01, P = 0.06). There was no longer a significant difference in CFU per lung between the WT and *pelA* mutant (P = 0.69). A significantly greater CFU per lung was recovered from the *gacA* mutant than the WT (P = 0.047) 7 d PI. (B) Micrograph of Gram-stained PA14 WT and selected transposon mutants in EVPL + ASM1 at 2 d and 7 d PI. All images are at x100 magnification.

We found that the standard crystal violet biofilm assay was not suitable for use on *ex vivo* tissue samples. As shown in Figure S6, the lung tissue binds large quantities of crystal violet obscuring any signal from dye bound to the biofilm matrix. H & E staining has been used in multiple studies and clinically as a quick, cost-effective approach to detect bacteria and their biofilms, which stain purple [53, 54]. Hence, we used H & E staining to visualise the *P. aeruginosa* biofilms on EVPL tissue. Replica sets of infected lung pieces from two independent lungs were fixed at 2 d PI and 7 d PI and stained with either Gram stain to confirm the bacterial species present (Figure 3B), or H & E to visualise the total biofilm mass (Figure 4).

**Figure 4.**
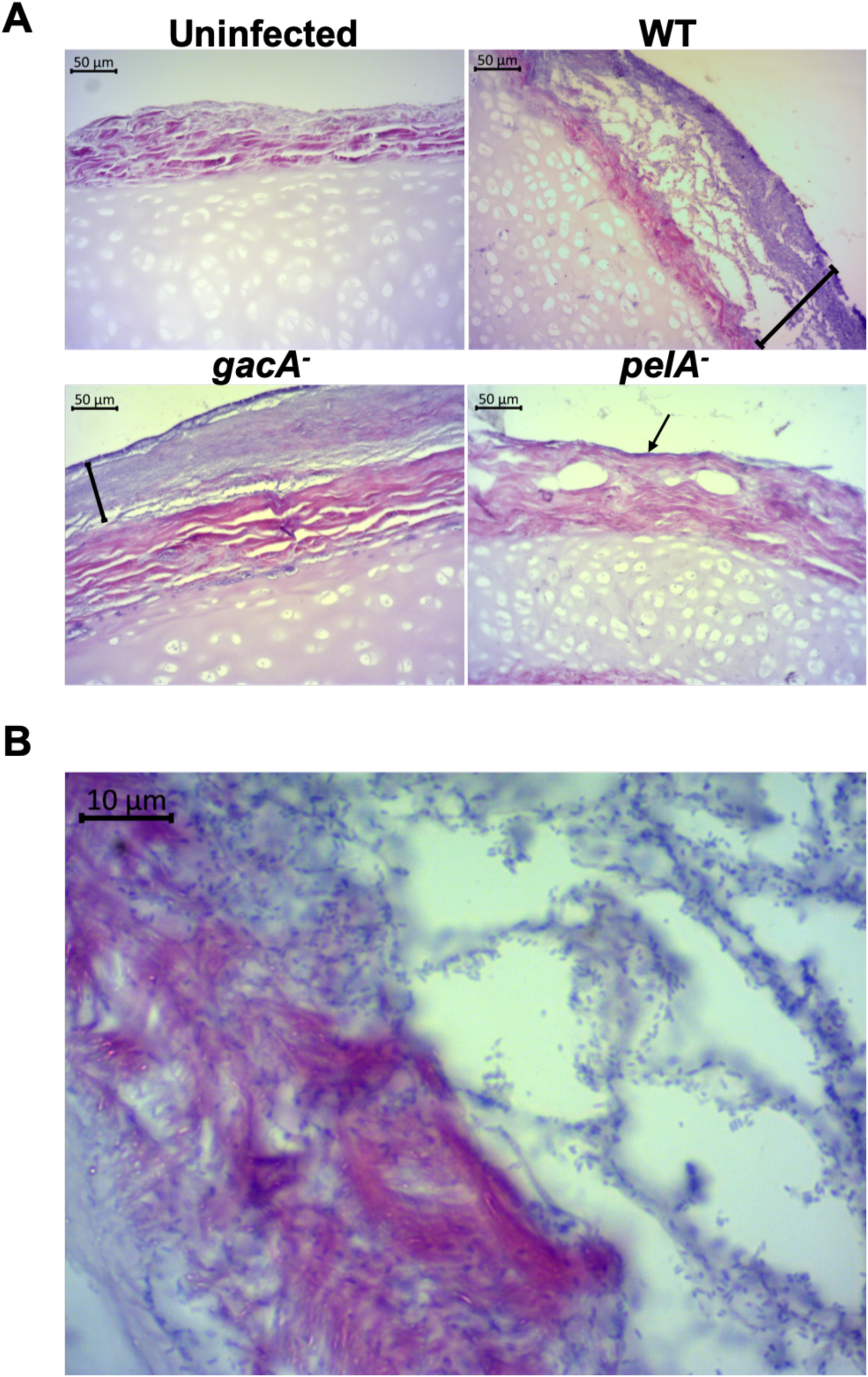
Hematoxylin & Eosin (H & E) stained sections of *ex vivo* pig lung (EVPL) bronchiolar tissue with artificial sputum media 1 (ASM1) at 7 d post infection (PI). The cartilage and tissue surface stain pink and the bacterial biofilm purple. (A) EVPL infected with *Pseudomonas aeruginosa* PA14 wild type (WT) and selected PA14 transposon mutants, with uninfected tissue as a negative control. All images are at x20 magnification. The presence of biofilm is denoted by black bars showing its depth for the WT and the *gacA* mutant, with an arrow for the *pelA* mutant (as the mutant forms only a very thin layer on the tissue surface). (B) PA14 WT infected EVPL at x100 magnification. The *P. aeruginosa* rods are visible, making up the structures evident in the lower magnification image in panel A.

Gram staining confirmed that the uninfected tissue was clear of bacterial growth at 2 d and 7 d PI (data not shown). Aggregates of Gram-negative rods were visible on the surface of EVPL tissue infected with each strain at both time points (Figure 3B). The Gram stain images also demonstrate the increase in density of bacterial cells from 2 d PI to 7 d PI for both mutants and the WT. Bacterial cells do not appear obviously stressed, maintaining their typical rod shape, in contrast with observations of *P. aeruginosa* grown in ASM1 *in vitro* (Figure S2).

H & E staining was also performed on replica sections from two independent lungs, and exemplar images for each strain at 7 d PI are shown in Figure 4 (replicate images from second lung: Figure S7). Although biofilm formation was detectable 2 d PI based on previously reported biofilm H & E staining [53, 54], the structure was not as defined (Figure S8). At both time-points the WT had formed a biofilm on the surface of the tissue. By 7 d PI, WT PA14 appeared to have formed a mature, structured biofilm (Figure 4A). We were also able to image individual WT cells stained with H & E. This showed their density and association with EVPL tissue 7 d PI, as well as a distinct lack of obvious stress over the prolonged infection period (Figure 4B).

There were consistent differences in the biofilm observed between the mutant strains and WT 7 d PI using H & E staining. This was also evident in the biofilm depth measurements taken from the images (Figure S9). At 7 d PI we observed a clear difference in the biofilm formed by the WT and *gacA* mutant (Figure 4A). The mature structure of the WT biofilm was not present in the biofilm formed by the *gacA* mutant: the purple stained region on the surface of the EVPL tissue demonstrates the homogenous, dense biofilm formed by the *gacA* mutant (Figure 4A). The *pelA* mutant exhibited greater observable differences in biofilm formation from the WT at both 2 d and 7 d PI (Figure S7; Figure 4B; Figure S9). Our images of the *pelA* mutant biofilm are comparable to the negative control uninfected tissue, despite presence of mutant infection found by Gram staining and CFU counts (Figure 3). We identified only a thin layer of purple stain at 7 d PI (Figure 4B). Although the *pelA* mutant was able to grow on EVPL tissue, it was unable to form a biofilm.

Our findings confirm that *P. aeruginosa* PA14 is able to form mature, highly structured biofilm communities on the surface of EVPL + ASM1. As expected for a chronic biofilm infection, both the Gac regulatory system and the Pel polysaccharide are required for this outcome.

## Discussion

CF has a major impact on not only an individual’s quality of life and high mortality rates, but also on healthcare systems worldwide. Throughout their lifetime CF patients continually require treatment for lung infections. In the US, the estimated median annual costs for treatment of pulmonary exacerbations, a period of acute-like infection that is highly virulent, ranges from $9,456 (USD) to $48,263 (USD) per patient [55]. There is a limited choice of effective antibiotics to treat CF lung infections currently available to clinicians, and the antibiotic tolerance observed *in vivo* means that standard antibiotic susceptibility testing does not accurately predict whether drugs will actually work [17, 18]. As CF patients are living longer, there is a growing medical and economic need for novel and improved treatments for lung infections that are more effective and less rigorous. The rising incidence of antimicrobial resistance (AMR) is only adding to this pressure. Prescribing cannot be improved, nor can a true insight into the impact of potential novel treatment agents (e.g. anti-virulence drugs) be achieved, until there is a better understanding of the fundamental microbiology underpinning these chronic biofilm infections. We aimed to develop a clinically-relevant *ex vivo* model for the predominant cause of chronic CF lung infections: *P. aeruginosa*. We have demonstrated that addition of EVPL tissue to a simple formulation of ASM results in a clinically realistic *P. aeruginosa* biofilm, overcoming a number of obstacles associated with current laboratory models of CF infection.

We initially aimed to define the optimal *P. aeruginosa* culture conditions for the EVPL model to ensure we are closely modelling clinical CF lung infections. Investigation of *P. aeruginosa* growth, virulence and purine biosynthesis showed that addition of the porcine bronchiolar tissue overcomes issues associated with *in vitro* growth in the laboratory media ASM1 and ASM2.

ASM1 is simpler and cheaper to make than ASM2, thus our conclusion that ASM1 is sufficiently realistic when used with EVPL makes the system readily accessible and tractable for use in research and potentially diagnostic laboratories. It is plausible that ASM1 is sufficient with the addition of EVPL for two primary reasons. Firstly, mucin is unlikely to be an important physiological cue *in vivo* and in EVPL as it is a poorly used carbon source for *P. aeruginosa* [49, 56]. *In vitro* it is likely that mucin acts as a structural cue for *P. aeruginosa* aggregation and the switch to biofilm lifestyle, however we observed biofilm formation on EVPL in the absence of exogenous mucin. Additionally, the prolonged growth periods and high biofilm densities possible with EVPL mean it is likely that membrane lipids, NAG and eDNA will be released from cells within the tissue and/or maturing biofilm. Our findings suggest that EVPL tissue with ASM1 provides key compounds that are lacking from the media alone, creating a closer match with authentic sputum from CF patients.

Visualisation of the biofilm is key to comparing laboratory models with *in vivo* biofilms. We found that the laboratory *P. aeruginosa* strain PA14 forms a highly structured biofilm on the surface of EVPL tissue. This mature biofilm architecture is comparable to typical *P. aeruginosa* biofilms that have been observed in explanted lung tissue from people with CF (see Figure 8 in [57], Figure 4 in [58] and Figure 4 in [16]). The EVPL biofilm appears more structured than has been previously observed *in vitro* [59], and distinct from the “mushroom” type typically seen in flow cell biofilms (see Figure 8 in [2]). Our findings support the idea that EVPL mimics a clinically realistic *P. aeruginosa* biofilm infection. The porcine tissue provides the structure and physiological cues that facilitate the formation of biofilms representative of CF infection.

To explore the mechanisms responsible for biofilm formation in the EVPL model, we also imaged the biofilm formed by two PA14 transposon mutants: *pelA* and *gacA*. We identified clear differences in growth in EVPL. The *pelA* gene encodes a deacetylase and hydrolase enzyme involved in the synthesis of the biofilm exopolysaccharide pellicle: Pel [52]. Previous studies have shown that the Pel polysaccharide is necessary for cell-cell interactions and development of *P. aeruginosa* PA14 biofilms [37]. This is consistent with our images showing that the *pelA* mutant was unable to form a biofilm on the surface of the EVPL bronchiolar tissue. The *gacA* mutant was found to form a biofilm, however it lacked the structure that was observed for the WT. GacA has been shown to be essential for *P. aeruginosa* PA14 biofilm maturation, and lack of *gacA* function results in a reduction in biofilm formation capacity and structure, and reduced antibiotic resistance [36]. Our observations thus confirm the importance of established biofilm regulatory pathways in the formation of the structured communities observed on EVPL tissue. This supports the conclusion that *P. aeruginosa* is forming a mature biofilm in the EVPL model.

Further analysis of EVPL infection at a genotypic level will be required to fully conclude which aspects of chronic CF infection dynamics the model is able to recapitulate. However, we present EVPL + ASM1 as a cheap, tractable and high-throughput platform for growing biofilms of *P. aeruginosa*. The EVPL infections we observed exhibit some of the key morphological and metabolic characteristics observed for this pathogen in CF. Our results clearly demonstrate the importance of visualising biofilms grown in laboratory models. When we compared the CFU per lung of WT, *pelA* and *gacA* mutants, there were no clear differences in bacterial recovery – but there were obvious differences in the structure of the biofilms they formed. Visualisation of biofilms formed on EVPL tissue will be a valuable addition to future work, with the aim to understand infection dynamics, antibiotic tolerance, polymicrobial interactions and the ability of novel drugs to penetrate and destroy these biofilms.

## Conclusions

We have shown, in extension to our previous work [30, 31], that the EVPL model is able to successfully replicate a phenotypically realistic *P. aeruginosa* CF lung biofilm. Our work can now be expanded to further understanding of other key aspects of infection, as well as a basis for development of *ex vivo* models for other biofilm infection contexts. The model provides a high-throughput approach to bridge the gap between *in vitro* models and true human infection.

## Acknowledgements

We thank all at Steve Quigley & Sons butchers for donating pig lungs; Leo Eberl, Steve Higgins, Sophie Darch, Steve Diggle and Paul Williams for bacterial strains; and John Moat and Ian Hands-Portman for help setting up imaging. The authors would also like to acknowledge the help of the Media Preparation Facility in the School of Life Sciences at the University of Warwick, with special thanks to Cerith Harries and Caroline Stewart. The Histology Facility equipment used in this study was purchased with grants from The University of Manchester Strategic Fund. Special thanks to Peter Walker and Grace Bako for their help with the histology. This work was supported by an MRC New Investigator Research Grant [grant number MR/R001898/1] awarded to FH and by a PhD studentship from the BBSRC Midlands Integrative Biosciences Training Partnership (MIBTP) awarded to NEH.

## Author Statement

**Niamh E. Harrington:** Conceptualization, Methodology, Validation, Formal analysis, Investigation, Writing – Original draft preparation, Visualisation.

**Esther Sweeney:** Methodology, Investigation, Writing – Review & Editing.

**Freya Harrison:** Conceptualization, Methodology, Validation, Formal analysis, Investigation, Writing – Review & Editing, Visualisation, Supervision, Project administration, Funding acquisition.

## Data Supplement

A data supplement will be provided with the final submission, containing all raw data and associated R code.

---

**Figure S1.**
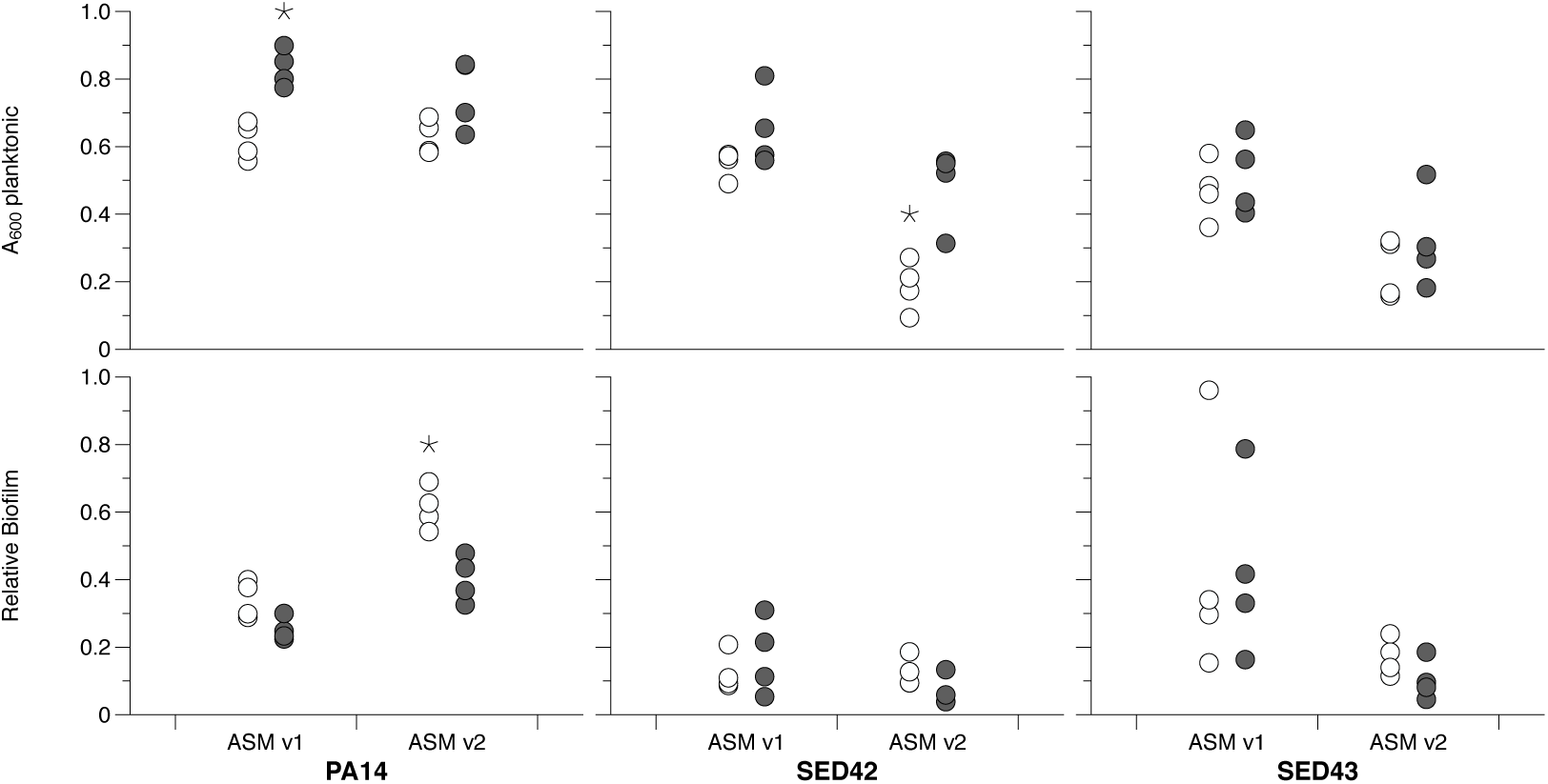
Growth of the *Pseudomonas aeruginosa* lab isolate PA14 and the Cystic Fibrosis (CF) isolates SED42 and SED43 in two versions of artificial sputum media (ASM) in 96-well plates with a peg lid to allow biofilm formation. Open circles: ASM with no glucose added. Closed circles: ASM supplemented with 3 µM glucose. Top panels show A_600mn_ of the planktonic subpopulations in each well after 24 h incubation at 37 °C. Bottom panels show relative allocation to biofilm on the plastic pegs, as measured by a crystal violet assay (A_592nm_ of crystal violet bound by each biofilm, divided by the A_600nm_ of the planktonic subpopulation of that well). Each data point is one replica population. ANOVA was used to analyse data and asterisks show conditions for each bacterial strain where growth or biofilm formation was significantly different when contrasted with that in ASM1 without glucose, using a post-hoc Tukey’s test. For PA14, planktonic growth was enhanced by addition of glucose (F_1,12_ = 23.7, *p* < 0.001) but was not affected by ASM version (main effect F_1,12_ = 0.73, *p* = 0.41; interaction with glucose F_1,12_ = 1.51, *p* = 0.24). Biofilm allocation by PA14 was higher in ASM2 than ASM1 (F_1,12_ = 56.0, *p* < 0.001) and reduced by the presence of glucose (main effect F_1,12_ = 28.7, *p* < 0.001; interaction with ASM F_1,12_ = 4.33, *p* = 0.059). For SED42, planktonic growth was lower in ASM2 than ASM1 (F_1,12_ = 33.0, *p* < 0.01) and enhanced by the addition of glucose (main effect F_1,12_ = 18.8, *p* < 0.01; interaction with ASM F_1,12_ = 4.69, *p* = 0.51). Biofilm allocation by SED42 was unaffected by either ASM version (F_1,12_ = 2.05, *p* = 0.18) or glucose (main effect F_1,12_ = 0.03, *p* = 0.86; interaction with ASM F_1,12_ = 2.06, *p* = 0.18). For SED43, growth was lower in ASM2 than ASM1 (F_1,12_ = 14.8, *p* = 0.002) and unaffected by glucose (main effect F_1,12_ = 1.17, *p* = 0.30; interaction with ASM F_1,12_ = 0.11, *p* = 0.74).

**Figure S2.**
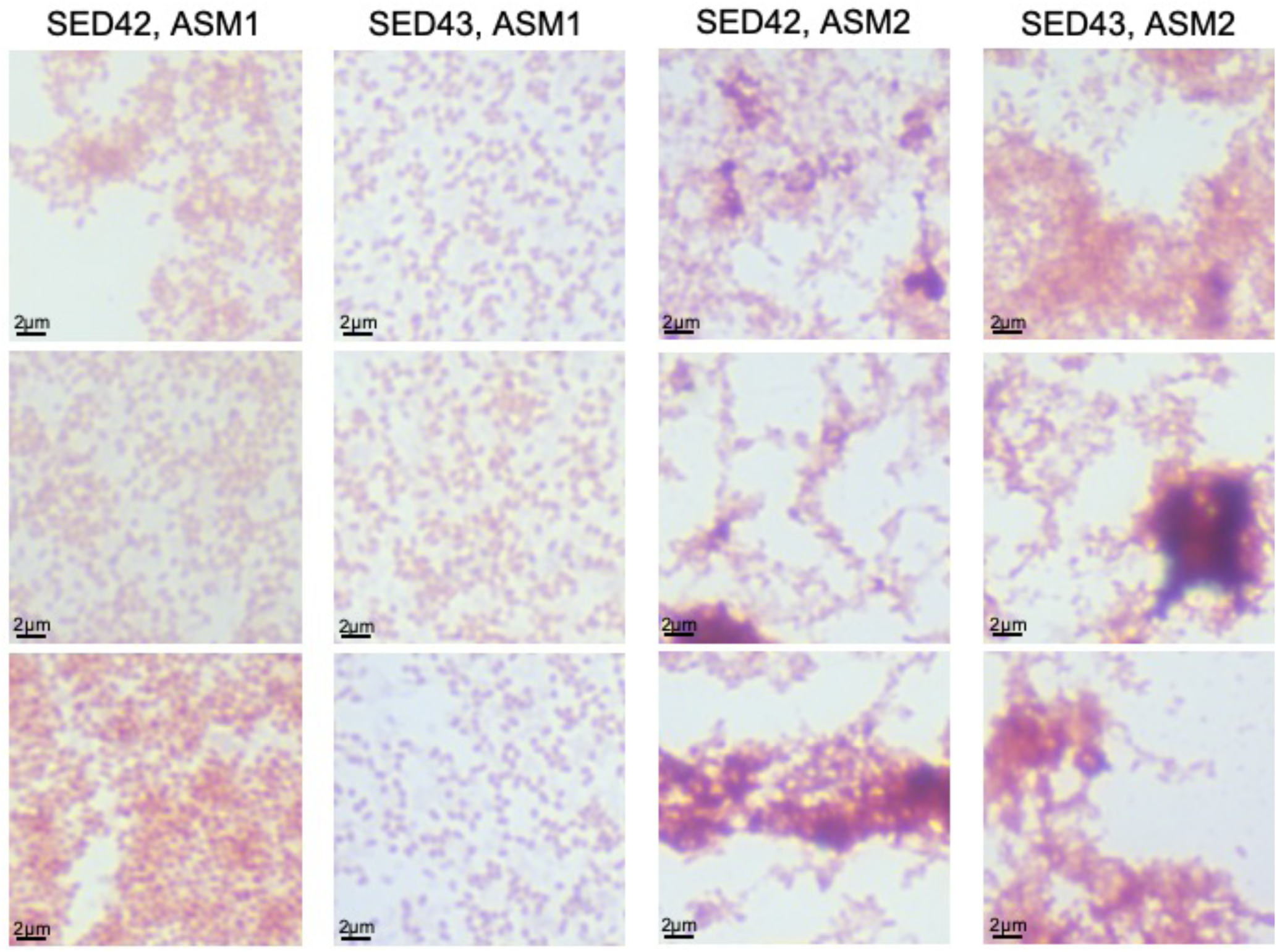
Example micrographs of Gram-stained *Pseudomonas aeruginosa* Cystic Fibrosis (CF) isolates after 36 h growth in two versions of artificial sputum media (ASM). Triplicate cultures were grown for each combination of strain and ASM version, and aliquots of culture heat fixed into a glass slide and Gram stained. Representative areas of approx. 20µm x 20µm are shown for each culture. All images are x100 magnification.

**Figure S3.**
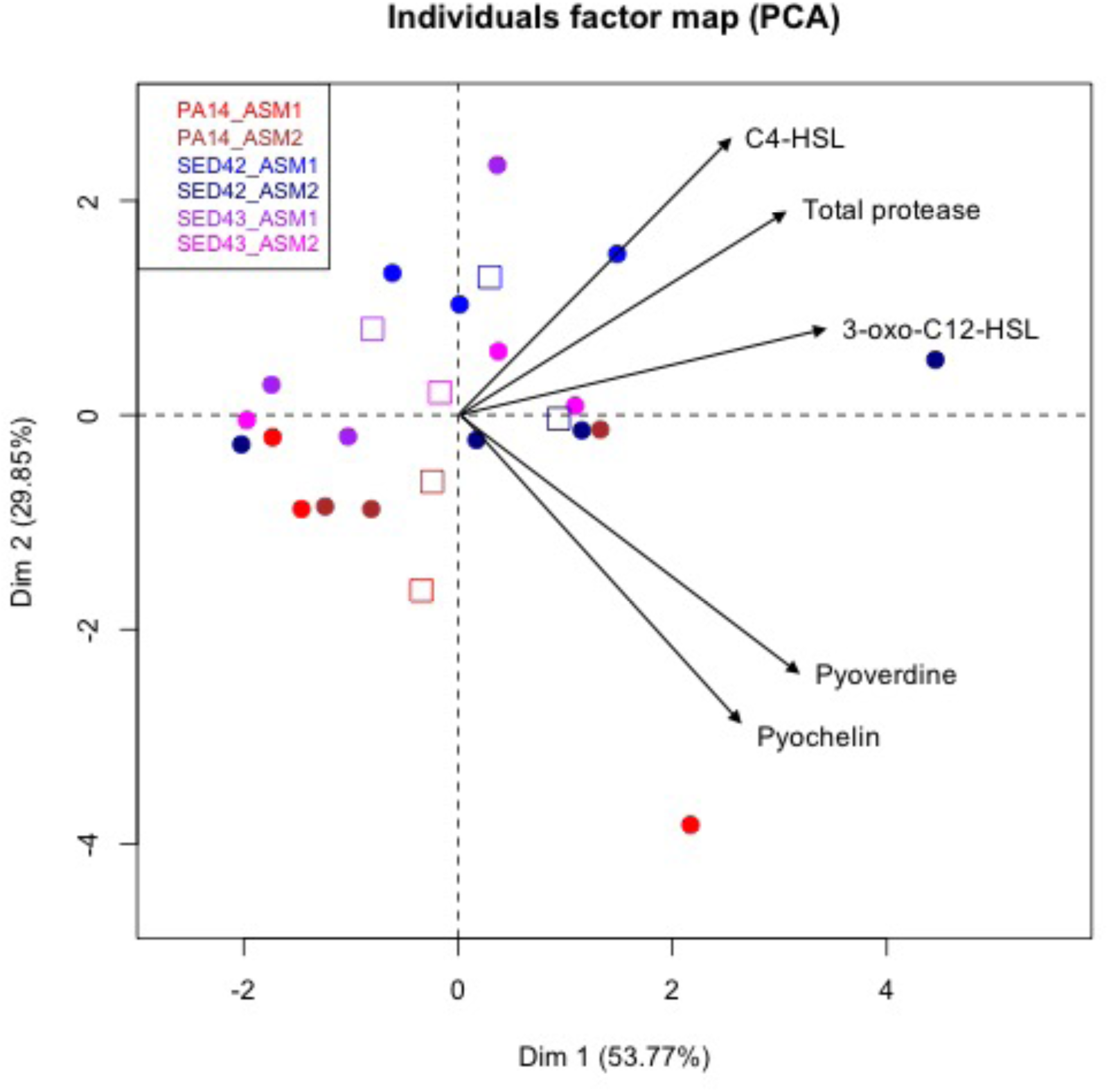
Principal component analysis of virulence factor data presented in Figure 1B for *Pseudomonas aeruginosa*. Only samples with data for all virulence factors were included. Circles show data points, squares are centroids. There is no clear clustering by artificial sputum media (ASM) version.

**Figure S4.**
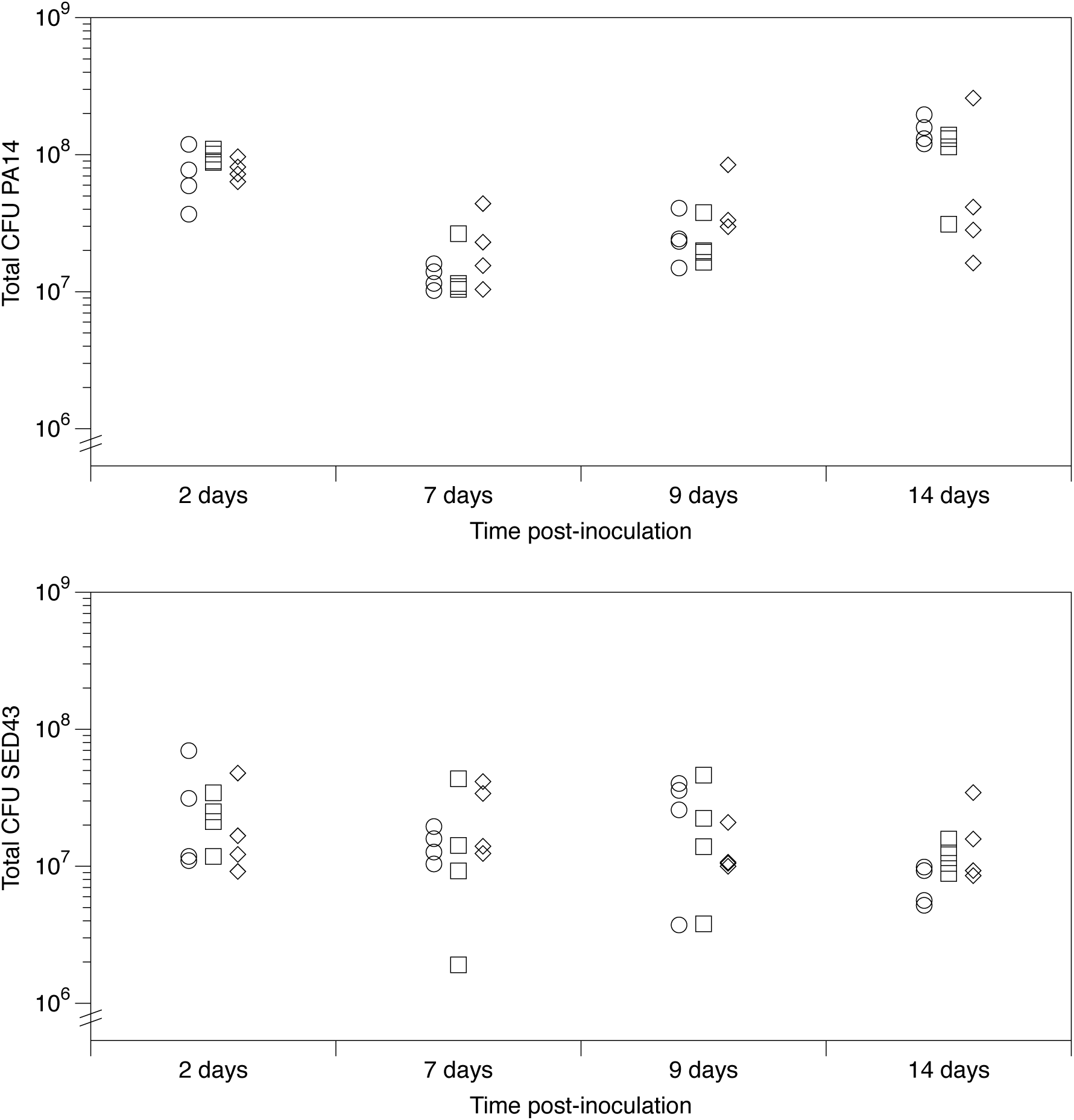
Growth of *Pseudomonas aeruginosa* PA14 and a representative chronic Cystic Fibrosis (CF) isolate: SED43, on pieces of *ex vivo* pig lung (EVPL) bronchiolar tissue plus artificial sputum media 1 (ASM1). Pieces of tissue from three independent lungs (different symbols) were inoculated with either PA14 or SED43 and destructively sampled at 2, 7, 9 or 14 d post infection (PI) at 37 °C to retrieve the biofilm. The graph shows colony-forming units (CFU) retrieved from individual EVPL biofilms at each time point. For PA14, after an approximate 1-log drop in CFU between 2 and 7 d PI, biofilm CFU increased again by 14 d, with some fluctuation between lungs. For SED43, CFU numbers were more consistent over time.

**Figure S5.**
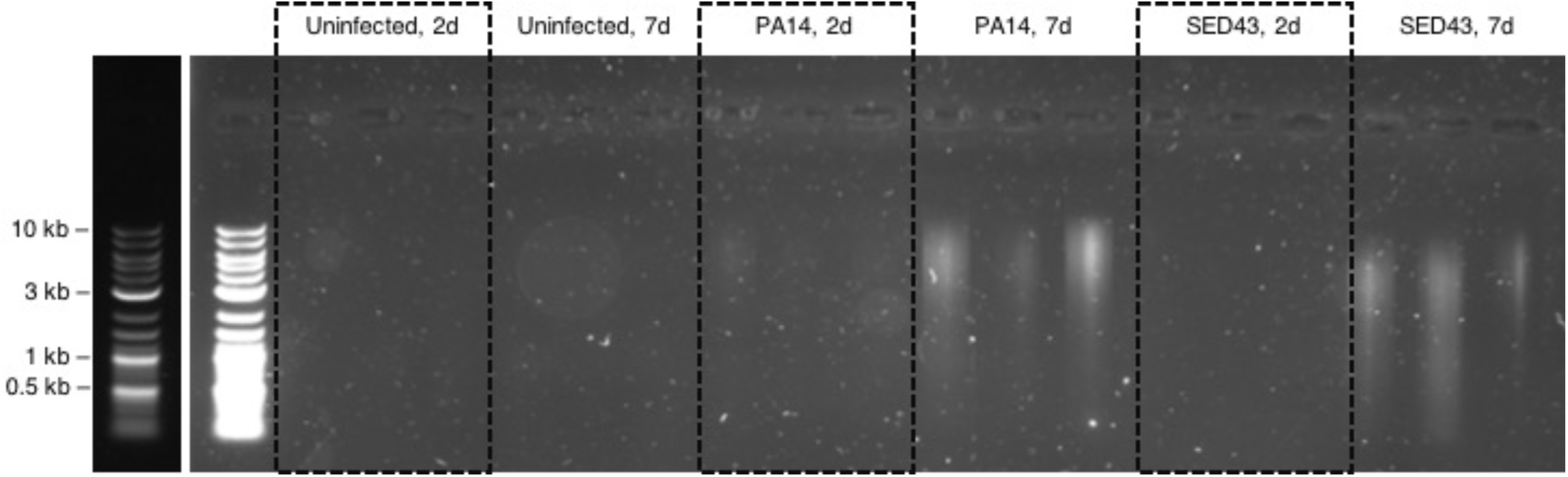
Detection of extracellular DNA (exDNA) in *Pseudomonas aeruginosa*-infected *ex vivo* pig lung (EVPL) + artificial sputum media 1 (ASM1). After 2 or 7 d growth, homogenate from EVPL cultures of PA14 and the clinical Cystic Fibrosis (CF) isolate SED43 (n=3 in each case) was centrifuged and the pellet run through a Promega Wizard Genomic DNA extraction kit. Recovered DNA was eluted in 50 µl water and 10 µl of the eluate run on an agarose gel. Uninoculated EVPL which had been incubated in ASM1 was also run through the extraction kit (n=3). No DNA was visible from uninoculated EVPL or EVPL after 2 d infection with *P. aeruginosa*, but after 7 d sufficient eDNA was present in *P. aeruginosa* biofilm to be visible on the gel. As the signal was so weak, the lane corresponding to the DNA ladder is shown in duplicate with different exposure times to facilitate contrast of the bands (NEB QuickLoad Purple 2-log ladder).

**Figure S6.**
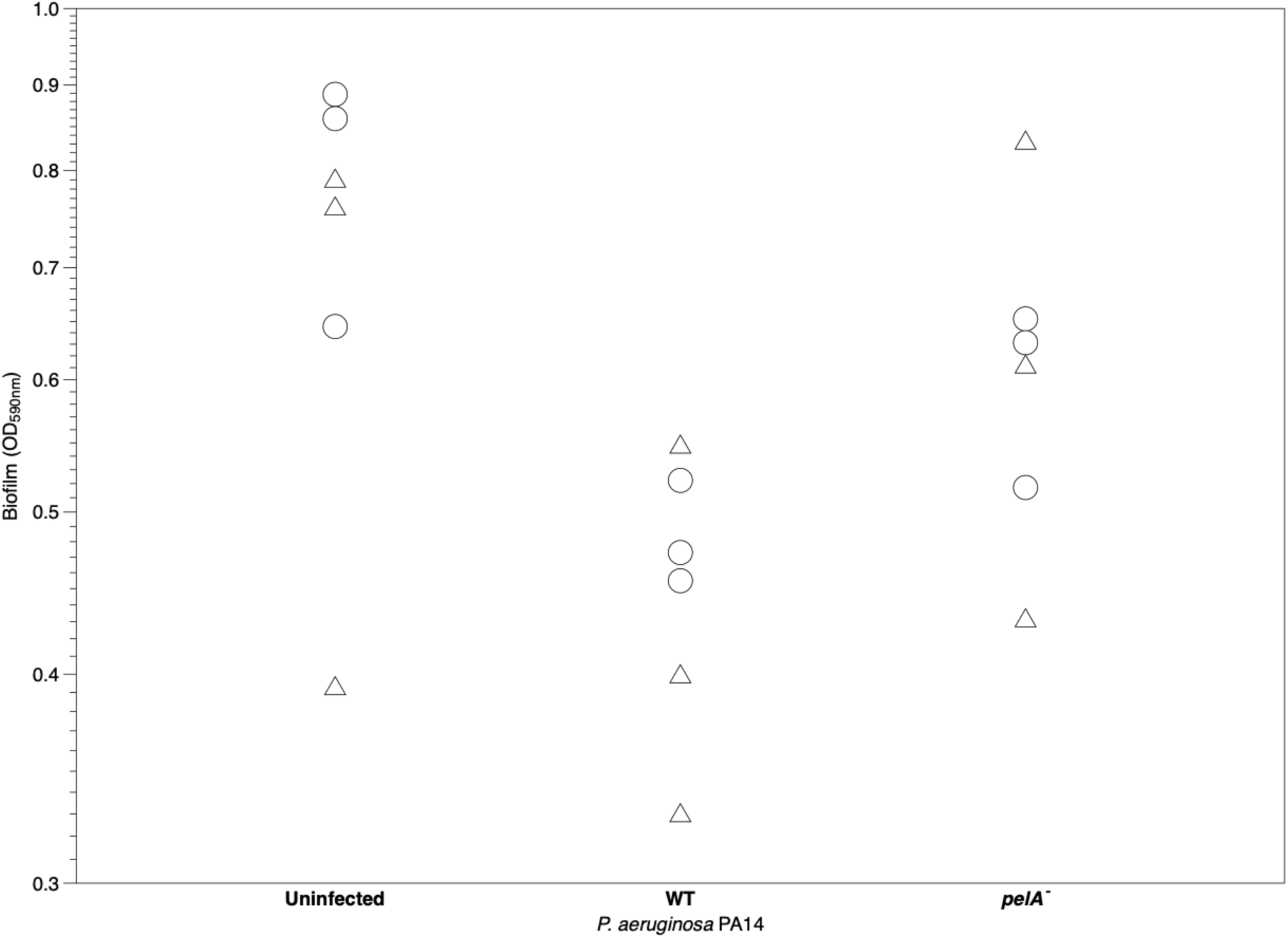
Measure of biofilm on *ex vivo* pig lung (EVPL) bronchiolar tissue with artificial sputum media 1 (ASM1) formed by *Pseudomonas aeruginosa* PA14 and a *pelA* transposon insertion mutant, from a crystal violet assay (A_590nm_ of crystal violet bound by each tissue sample). Three replicate pieces from two independent lungs were infected with each strain. The EVPL tissue was infected for 2 d at 37 °C. Uninfected tissue was used as a negative control. The tissue pieces were washed in PBS and repeatedly stained with 0.1% (v/v) crystal violet. Following drying of stained tissue, the bound crystal violet was solubilised in 95% (v/v) ethanol and the absorbance read at 590 nm. The highest absorbance measured was for the uninfected tissue sample, indicating that the crystal violet preferentially binds to the tissue over the biofilm. Therefore, this assay was considered unsuitable to measure EVPL-formed biofilms.

**Figure S7.**
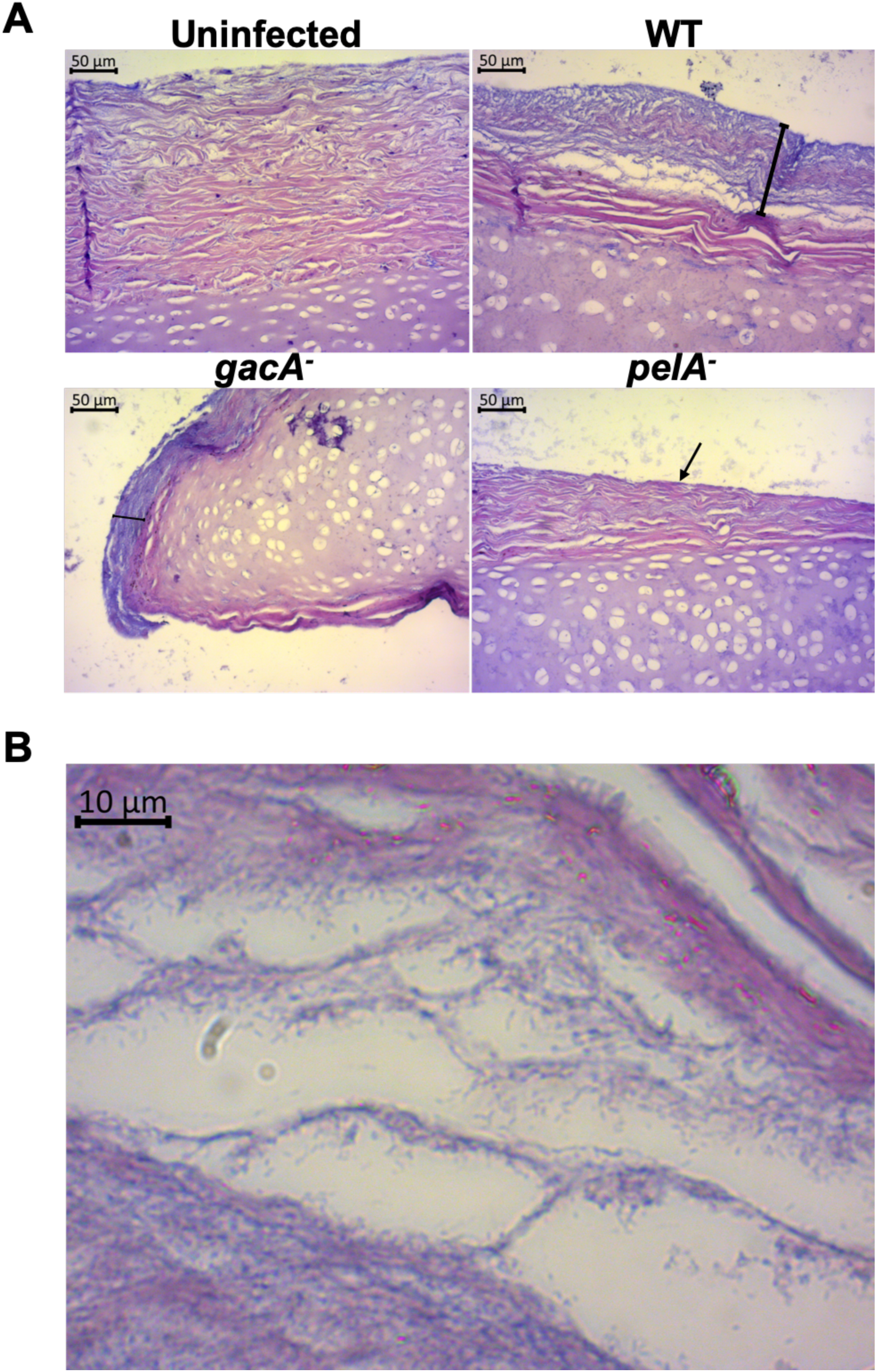
Hematoxylin & Eosin (H & E) stained sections of *ex vivo* pig lung (EVPL) bronchiolar tissue with artificial sputum media 1 (ASM1) at 7 d post infection (PI). Two independent lungs were infected for 7 d, fixed and H & E stained. Figure 4 images were taken from one lung and these images were taken from a second lung. The cartilage and tissue surface are stained pink and the bacterial biofilm purple. (A) EVPL infected with *Pseudomonas aeruginosa* PA14 WT and selected PA14 transposon mutants, with uninfected tissue as a negative control. All images are at x20 magnification. The presence of biofilm is denoted by black bars showing its depth for the WT and the *gacA* mutant, with an arrow for the *pelA* mutant (as the mutant forms only a very thin layer on the tissue surface). (B) PA14 WT infected EVPL at x100 magnification. The *P. aeruginosa* rods are visible, making up the structures evident in the lower magnification image in panel A.

**Figure S8.**
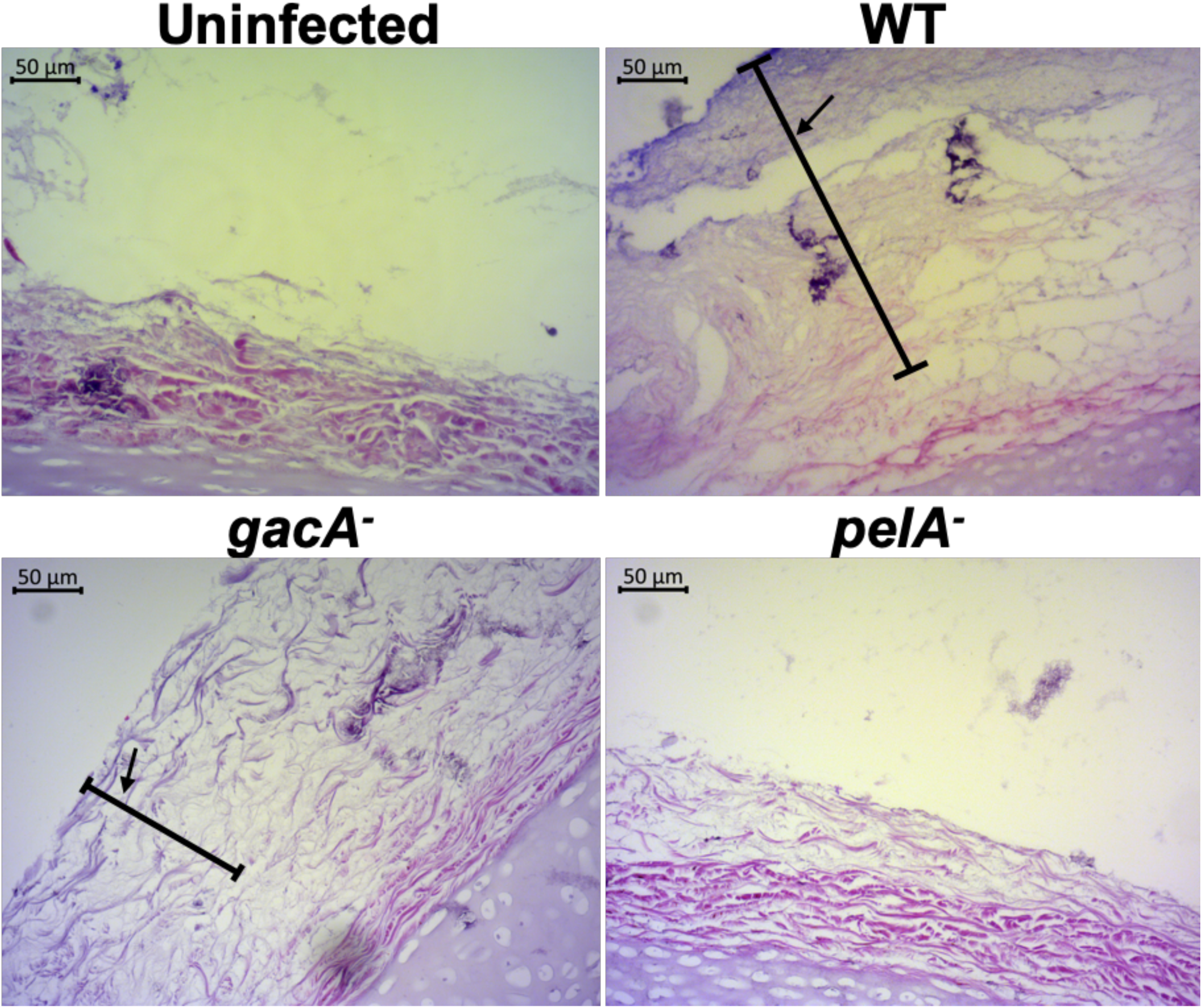
Representative micrographs of *ex vivo* pig lung (EVPL) bronchiolar tissue with artificial sputum media 1 (ASM1) 2 d post infection. EVPL was individually infected with wild type (WT) *P. aeruginosa* PA14 and selected transposon mutants for the PA14 strain. Uninfected tissue was used as a negative control. All tissue samples were paraffin embedded then stained with Hematoxylin and Eosin (H & E). All images are at x20 magnification. The tissue surface is stained pink and the bacterial biofilm in purple. The arrows and bars indicate the stained biofilm on the surface of the tissue.

**Figure S9.**
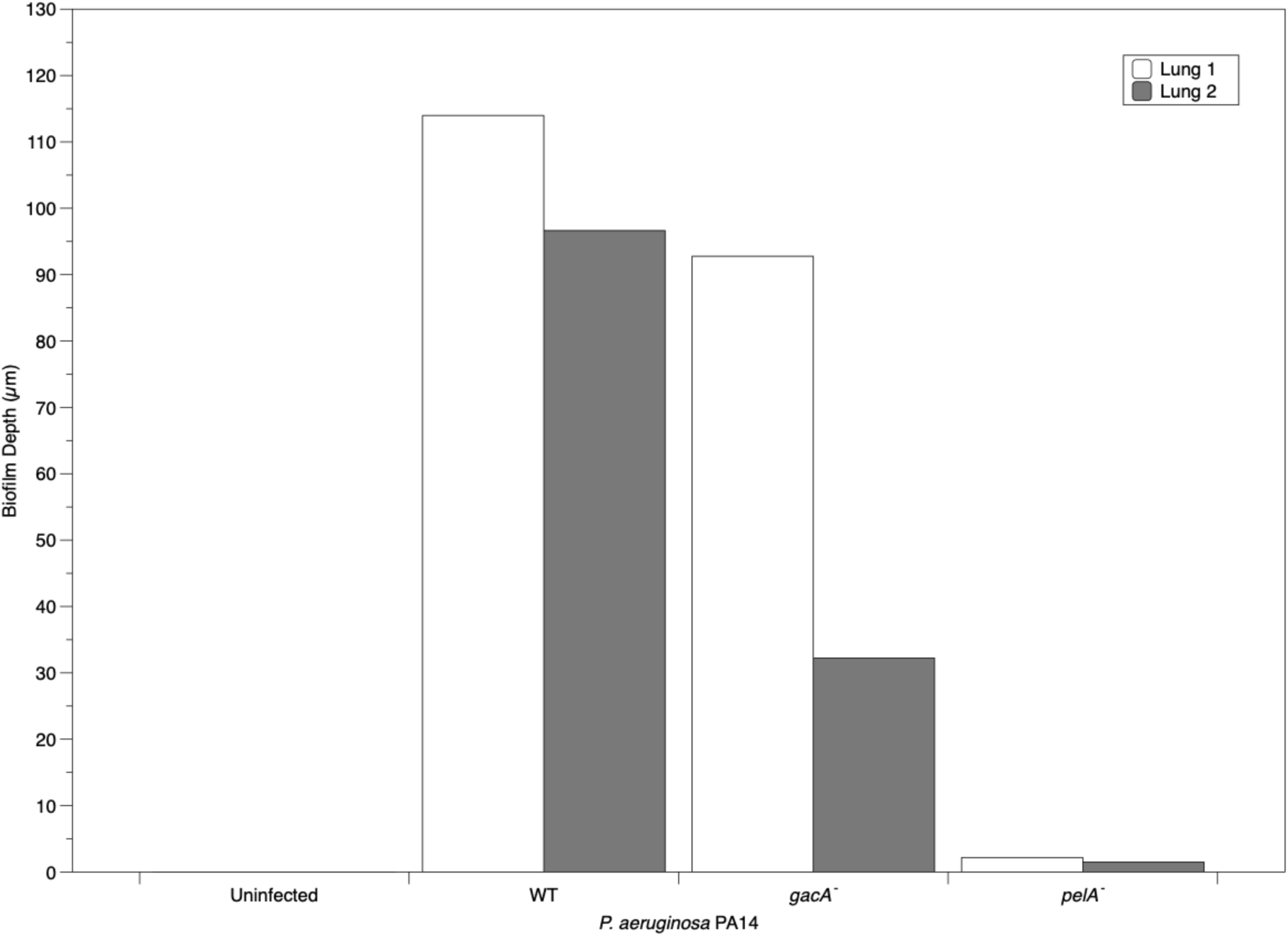
The depth (µm) of *Pseudomonas aeruginosa* biofilms formed on the surface of *ex vivo* pig lung (EVPL) bronchiolar tissue with artificial sputum media 1 (ASM1) 7 d post infection. Following infection of tissue pieces from two independent lungs with isogenic wild type (WT) *P. aeruginosa* PA14 and selected PA14 transposon mutants - uninfected tissue used as a negative control - tissue samples were paraffin embedded and Hematoxylin and Eosin (H & E) stained. Four measurements were taken along the biofilm from each H & E image using Zeiss Zen 2.3 pro software. The average biofilm depth per WT, mutant and uninfected tissue in the two lung repeats is shown.

